# Divergent spike mutations impact the activation of the fusion core in Delta and Omicron variants of SARS-CoV-2

**DOI:** 10.1101/2023.10.19.563184

**Authors:** Mandira Dutta, Gregory A. Voth

**Affiliations:** Department of Chemistry, The University of Chicago, Chicago, IL 60637; Chicago Center for Theoretical Chemistry, The University of Chicago, Chicago, IL 60637; Institute for Biophysical Dynamics, The University of Chicago, Chicago, IL 60637; James Franck Institute, The University of Chicago, Chicago, IL 60637

## Abstract

SARS-CoV-2 infects host cells by binding the receptor-binding domain (RBD) of its spike protein to the receptor, ACE2. A subset of highly effective spike mutations plays critical roles in altering the conformational dynamics of spike protein. Here, we use molecular dynamics simulations to investigate how spike mutations affect the conformational dynamics of spike/ACE2 complex in the D614G, Delta (B.1.617.2) and Omicron (B.1.1.529) SARS-CoV-2 variants. We observe that the increased positive-charged mutations in the Omicron spike amplify its structural rigidity and reduce its structural flexibility. The mutations (P681R in Delta and P681H in Omicron) at the S1/S2 junction facilitate S1/S2 cleavage and aid the activation of the fusion core. We report that high structural flexibility in Delta lowers the barrier for the activation of the S2 core; however, high structural rigidity in Omicron enhances the barrier for the same. Our results also explain why Omicron requires the presence of a higher number of ACE2 to activate its fusion core than Delta.

## Introduction

Despite the extensive development and improvement of effective vaccines for severe acute respiratory syndrome coronavirus 2 (SARS-CoV-2), the emergence of deleterious mutations in the virus is preventing the return to the pre-pandemic conditions so far (which some predictions suggest will never happen). These circumstances motivate the need to improve our knowledge of the role of mutations that impact virus adaption and fitness. Some studies have reported that the highly-effective mutations associated with virus phenotype in regards to their fitness are a few in number compared to ’low-effect’ or ’no-effect’ mutations(*1*). The advent of the first variant (Wuhan-Hu-1) was reported in the city of Wuhan, China, in December 2019(*2, 3*), followed by a series of variants with increased mutations, including the Alpha, Beta, Kappa, Delta, and Omicron variants, and the number is still rising at an alarming rate(*4*). As of December 2022, three major waves of Covid-19 infection that were believed to be associated with the D614G mutated variant of Wuhan-Hu-1 (named D614G from now on in this article), the Delta, and the Omicron caused a death toll of millions of people worldwide(*5–7*). Delta is reported to be six-fold less sensitive to serum-neutralizing antibodies and eight-fold less sensitive to vaccine-elicited antibodies compared to D614G(*8*). In November 2021, the Omicron variant was first reported in South Africa and spread quickly in numerous countries due to its higher infectivity and transmissivity(*6*). Although the emerging variants often outcompete the existing variants, each of them still builds steps that may support other future variants. Our present study focuses on Delta (B.1.617.2) and Omicron (B.1.1.529), including D614G, which have significantly impacted the SARS-CoV-2 pandemic history so far. Understanding the critical roles of the mutations in these variants will be helpful in controlling the pandemic situation and for the improvement of vaccines.

This virus is composed of four major structural proteins, including spike (S), nucleocapsid (N), envelope (E), and membrane (M)(*9, 10*). The S protein is a heavily glycosylated trimeric membrane protein that anchors the viral membrane with its transmembrane domains(*9, 11, 12*). It first binds to the host cell receptor, angiotensin-converting enzyme 2 (ACE2), and undergoes large conformational rearrangements to mediate the fusion between the viral and host cell membranes(*9, 13*). A full-length S monomer is divided into two sub-domains (Figure 1), S1 and S2. S1 comprises an N-terminal domain (NTD), a receptor binding domain (RBD), and a C-terminal domain (CTD). On the other hand, the S2 domain encodes a fusion peptide (FP), a fusion-peptide proximal region (FPPR), a heptad repeat 1 (HR1), a central helix (CH), a connector domain (CD), and a transmembrane segment (TM)(*14*). All these variants are associated with several mutations in the spike domain. Zhang *et al.* resolved the structures of the spike proteins of Delta(*15*) and Omicron variants using cryo-electron microscopy (cryo-EM)(*16*). There are six mutations in the spike of Delta and thirty-two mutations in the spike of Omicron, including D614G (Table S1 of the Supporting Information). Several studies reported that a higher number of charged mutations in the RBD domain of Omicron makes it more efficient to bind with the ACE2 receptor(*17–20*). Recently, Dokainish *et al*. observed that mutated D614G spike losses the interaction between D614 (S1) and K854 (S2) that stabilizes the FPPR loop and the furin cleavage site and provides an explanation for the higher stability for the pre-fusion state in D614G structure(*21*). They observed that this particular mutation makes the 630-loop ordered, which allosterically reorganizes RBD for ACE2 binding. A population shift was observed towards the one-state up open RBD conformation with D614G structure, suggesting a potential explanation for the enhanced infectivity of the variant(*22*). P681 residue is located at the furin cleavage site, separating S1 and S2 domains. The reverse mutation of R681 residue to PRO in Delta significantly reduces the replication than the alpha variant(*23*). An improvement in spike cleavage was also reported in the presence of P681R mutation in Delta. Casalino *et al.* studied the role of spike glycans in closed-to-open transition in RBD(*24*). They specified two N-glycans at sites N165 and N234, which play critical roles in regulating the conformational dynamics of RBD. Another study by the same group also revealed a gating mechanism of N-glycan at position N343, which modulates RBD opening(*25*). Hsiao *et al*. highlighted the critical role of glycans at the RBD/ACE2 interface of Omicron and the interactions between glycans and mutated residues results in enhanced binding with receptor(*26*). The “bottom-up” multiscale coarse-grained studies with D614G spike protein revealed that S protein binding to ACE-2 likely leads to complete dissociation of the S1 domain and unmasks the S2 fusion core(*27*). The cooperative interactions involving several ACE2 bound to the same S protein trimer are required to shed the S1 domain and activate the S2 core for viral fusion(*27*). Interestingly, Meng *et al*. reported that Omicron has lower replication in lung organoids and epithelial cells than Delta and D614G(*28*). Additional studies also reported sufficiently lower viral load in Omicron-infected lung epithelial cells compared to the Delta and D614G(*28–30*). In contrast, an increase in viral replication was observed in the Omicron-infected nasal epithelial cells. Moreover, Omicron exhibits lower replication and fusion activity than Delta in TMPRSS2-expressed cells(*31*). All of these observations hint at mechanisms that could be a potential reason for the higher transmissibility and reduced severity of Omicron over Delta SARS-CoV-2. As such, further investigations are needed to explore the underlying mechanisms.

**Figure 1.**
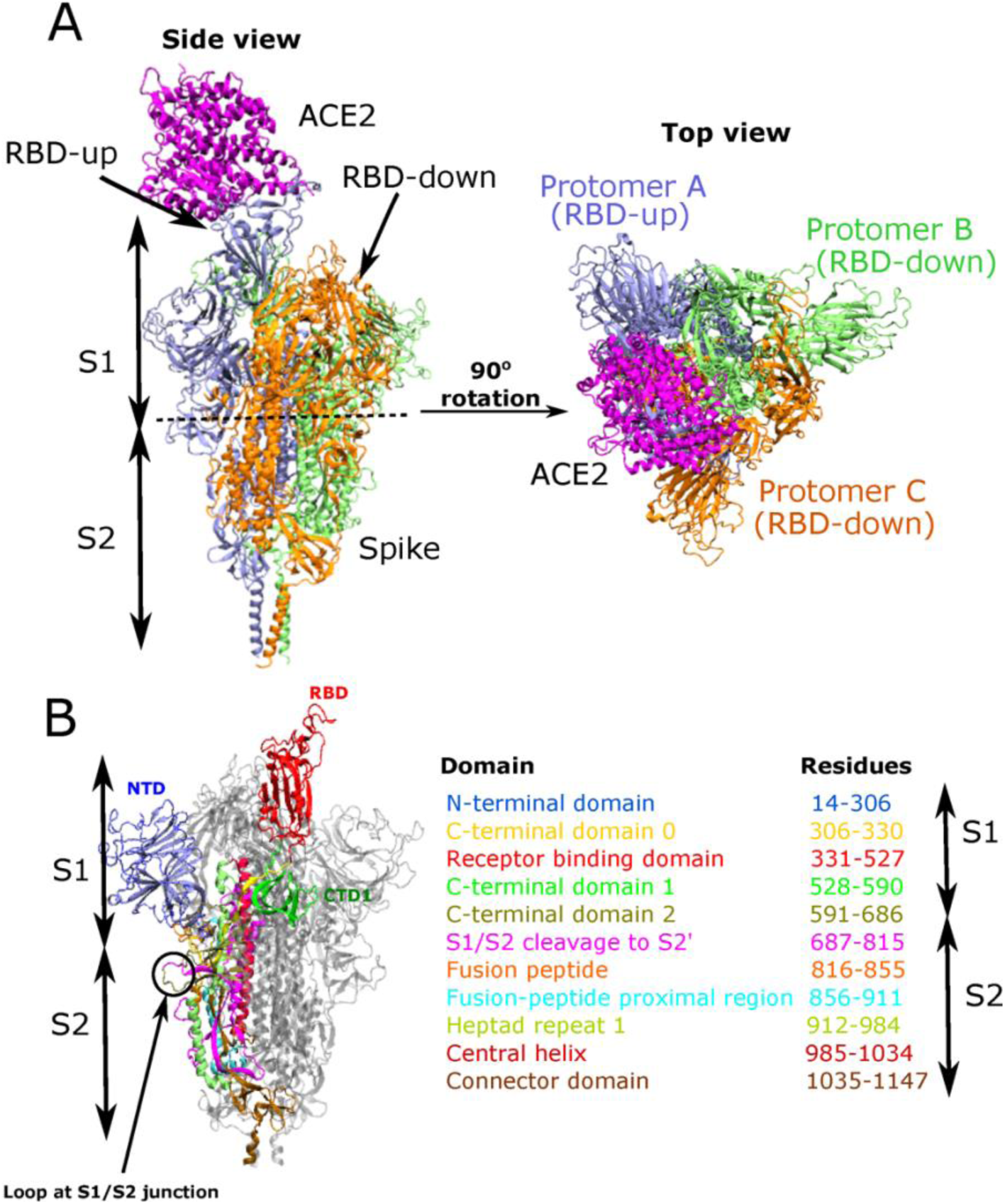
Structure of ACE2 bound SARS-CoV-2 spike protein. A) Side (left) and top (right) views of trimeric spike-protein/ACE2 complex structure. The blue, green, and orange color are the three protomers of the spike. ACE2 is shown in magenta. Protomer-A has RBD-up states, and protomer- B and C have RBD-down states. B) Different subdomains of spike protein are labeled in different colors. The S1 domain ranges from 14 to 685, and the S2 domain ranges from 686 to 1147.

To illustrate the impacts of mutations on spike protein dynamics, we performed molecular dynamics (MD) simulations of glycosylated spike proteins of D614G, Delta, and Omicron in their ACE2-bound complex. We observed that ACE2 binding initiates swing motions at the RBD domain that escalates the thermal fluctuation of the S1 domain. At the S1/S2 junction, the P681 residue is mutated to the considerably larger ARG residue for Delta and PHE for Omicron variants. These mutations assist in the partial unfolding of the loop, which leads to an escalation in S1/S2 cleavage and an increase in the fluctuation of FPPR regions, facilitating the distribution of the thermal motions from the RBD to the S2 fusion core. This, in turn, aids the large and complex conformational changes at the HR1/CH junction during pre- to post-fusion transition. However, for Omicron, the thermal fluctuation cannot compensate for the stronger intra and inter-domain interactions coming from a large number of positively charged mutated residues. Zhang et al. reported that Omicron S protein shows slightly less cleavage between S1 and S2 at 24h posttransfection, suggesting N679K and P681H mutations at the furin cleavage site do not assist S protein processing(*16*). In Delta, we observed a slight swelling of the S2 core. The free energy profile calculated over backbone torsion time-lagged independent component analysis (tICA) of the linker between HR1 and CH junction shows two conformational samplings, i.e., bent and slightly stretched structures. However, a single minimum with a bent conformation of the linker is obtained for both D614G and Omicron (Figure 7). Our results suggest that due to the higher number of mutations and less flexibility of the Omicron spike, it requires the presence of a higher number of ACE2 to activate its fusion core.

We note that the human lung has lower ACE2 than the nasal pathway(*32, 33*). Therefore, we provide here a possible explanation of why Delta shows higher viral load in the lung cells while Omicron preferably stays in the nasal tract. Our study is therefore essential to understand the molecular mechanisms of the viral membrane fusion and the critical roles of the spike mutations, starting from the receptor binding at RBD to the activation of S2 fusion machinery. This work may also aid further research in targeted and modifying existing drugs and vaccines.

## Results and Discussion

In our current study, we performed 3 *μs* all-atom MD simulations of the trimeric spike protein of D614G, Delta, and Omicron variants in complex with ACE2 (Figure 1A). We simulated two replicas for each system. We did not take explicit membrane in our simulation; however, to consider the effect of the membrane, we restrained the positions of the backbone atoms at the end of the TM (residue 1150-1162) segment. The ACE2-bound trimeric spike protein in these simulations was fully glycosylated and had one RBD-up (protomer-A) and two RBD-down protomers (protomer-B and protomer-C).

### Modulation of RBD/ACE2 swing motions in the presence of spike mutations in Delta and Omicron

We observed that ACE2 binding initiates swing motions in the RBD domain. To quantify the swing motions, we calculated the principal components (PCs) of ACE2 bound RBD movements (Figure 2). The first three principal components cover more than 40% of the overall motion of Omicron and more than 50% and 60% of the overall motions of D614G and Delta, respectively (Figure 2A). Figure 2B shows the first three principal motions of the RBD-ACE2 structure. Eigenvector-1 corresponds to a swing motion of ACE2 bound RBD approaching and leaving its neighbor down RBD. Eigenvector-2 describes a swing motion of ACE2-RBD towards its down and up states. Eigenvector-3 implies a motion toward the neighboring NTD domains. The Supporting Information (SI) contains the movies of the first three motions of the RBD-ACE2 structure (movie S1, movie S2, and movie S3, respectively). These swing motions escalate the thermal fluctuations of the S1 domain. Figure S1A of SI represents the projection of PC1 vs. PC2, which was calculated based on the coordinates of the C*α* atoms. We observed a larger spread of the eigenvector-1 and eigenvector-2 for Delta, followed by D614G. Omicron exhibits a confined spreading of the eigenvectors. The results suggest Delta visits more conformational space due to its structural flexibility than Omicron. A cryo-EM experiment revealed the ACE2 bound conformation of trimeric S glycoprotein and further reported that ACE2 binding induces continuous swing motions in RBD, which propagate to the neighboring RBD domains and perturb the allosteric network of the fusion machinery(*34*). Raghuvamsi *et al*., in their amide hydrogen–deuterium exchange mass spectrometry experiment, noted that ACE2 binding at the RBD exhibits large-scale changes in deuterium exchange in S1/S2 cleavage site and HR1 domains, which implies ACE2 binding at the RBD allosterically propagates to the distal sites of the S protein(*35*). We hypothesize that spike mutations play critical roles in altering the propagation of allosteric communication from the RBD to the fusion core.

**Figure 2.**
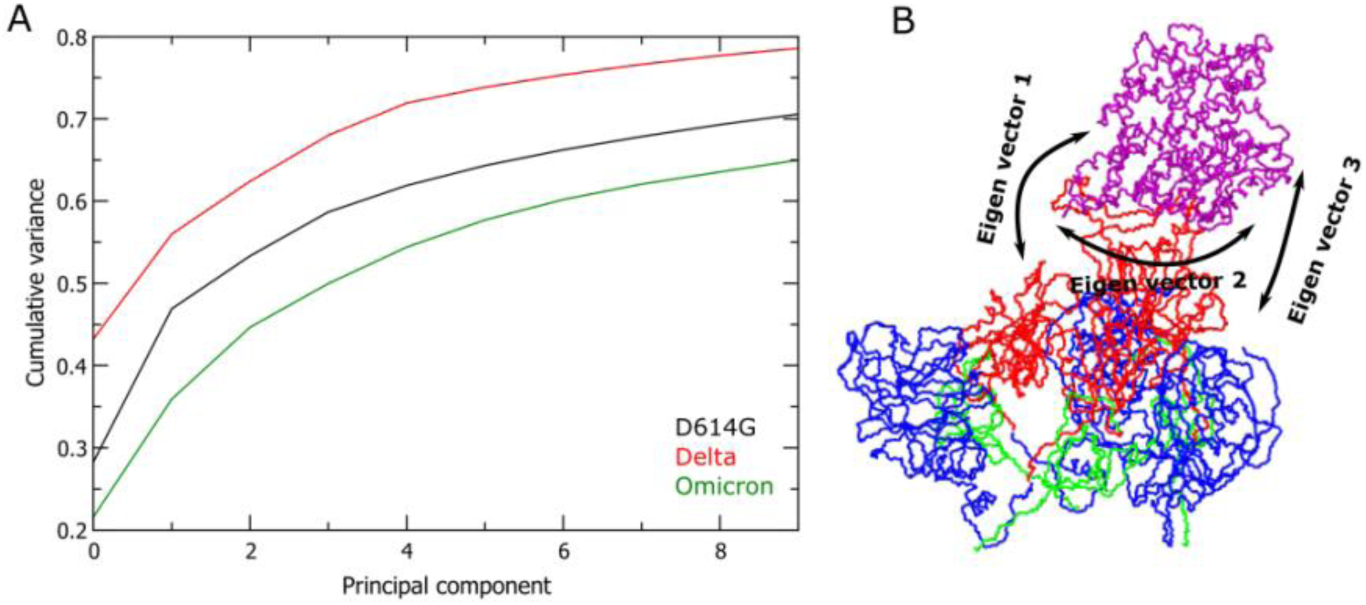
Principal component analysis (PCA) of RBD/ACE2 bound spike of SARS CoV-2 variants. Panel A) is principal component vs. cumulative variance for SARS CoV-2 ACE2/S1 complex. Black, red, and green colors show D614G, Delta, and Omicron variants, respectively. Panel B) It shows the first three principal motions described as eigenvector 1, eigenvector 2, and eigenvector 3. Blue, red, and green colors represent the NTD, RBD, and CTD domains of S1, respectively, and magenta represents ACE2.

### Spike mutations in Omicron enhance structural rigidity in S1 and S2 domains compared to Delta

To verify our hypothesis, we looked into the surface charge distribution and root mean square fluctuation (RMSF) of the RBD-up protomer to help explain the flexibility of the structure. Figure 3A, B represent the electrostatic surface plots of protomer-A of Delta and Omicron. We observed the presence of a high number of positive charges (indicated by the blue color) in both the S1 and S2 surfaces of Omicron, resulting in reduced conformational fluctuations and a relatively rigid structure compared to Delta. To investigate the structural flexibility, we determined the RMSF of the spike residues of protomer-A (Figure 3C, D, E) and protomer-B and C (Figure S2A, B of SI). For more clarity, we divided the RMSF plots in Figure 3 into three distinct regions of lower, medium, and higher fluctuations based on three cut-offs. For the lower fluctuation zone, we considered up to 0.2 nm (pink color), for medium and higher fluctuations, we used boundaries of 0.2 nm to 0.3 nm and above 0.3 nm (orange and green), respectively. We found that most of the Omicron S residues fall into the lower fluctuation area, whereas most Delta S residues are distributed between medium to high areas. D614G variant has more residues in the higher fluctuation zone than Omicron. However, the numbers are slightly lower, especially in the S2 domain, compared to Delta.

**Figure 3.**
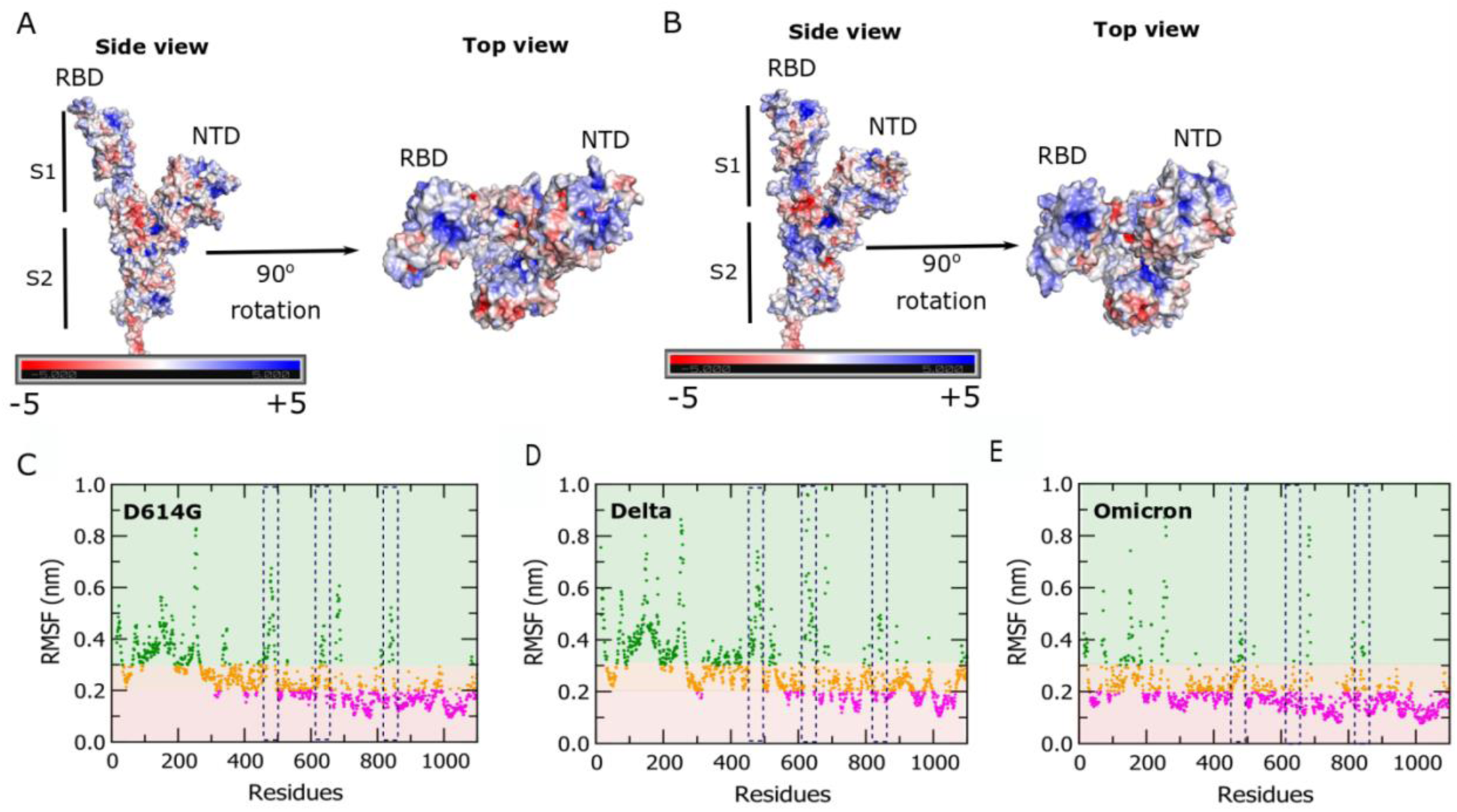
Electrostatic surface plots and RMSF plots of SARS CoV-2 spike protein. Panels (A) and (B) show electrostatic surface plots of spike protomer-A of Delta and Omicron, respectively, in two different orientations (side view and top view). The red-to-blue color bar indicates negative to positive charge distribution. Panels (C), (D) and (E) show RMSF plots of spike protomer-A of D614G, Delta, and Omicron variants. The RMSF values are separated into three distinct regions of lower, medium, and higher fluctuation zones. Pink, which ranges from 0 to 0.2 nm, defines lower fluctuation. Orange and green show residues in the medium (0.2 nm to 0.3 nm) and higher (above 0.3 nm) fluctuation regions. Three regions are highlighted in dotted lines. From left to right, they are RBM, 630-loop, and FPPR.

We have also highlighted three regions through dotted lines in RMSF plots. The first one from the left side is the receptor binding motif (RBM) (residue 468-490), which directly interacts with the ACE2 receptor. It shows lower fluctuation in Omicron than Delta and D614G, suggesting Omicron has stronger binding with the receptor than the other two variants. The middle and last ones are the 630-loop (residue: 620-640) and FPPR (residue: 828-853), which are essential components of S fusion machinery(*36–38*). Both remain structured in the RBD-down conformation. The 630-loop stays buried into a gap between the NTD and CTD1, stabilizing CTD2 of the same protomer when the RBDs are down. During the RBD opening, both 630-loop and FPPR move out of their positions, which enhances their structural flexibility and causes the loops to disorder(*39*).

We observed higher fluctuations for both the 630-loop and FPPR in Delta than in Omicron. For RBD-down protomers (Figure S2 of SI), we mostly noticed lower fluctuation values of the residues in D614G and Omicron, whereas Delta showed increased RMSF values in the S1 domain. The large swing motions of the ACE2 bound RBD domain of Delta spread to the neighboring RBD-down protomers that enhance the fluctuations of the S1 domain of protomer-B and C. We also calculated the solvent accessible surface area (SASA) for the 630-loop to examine its outward movement (Figure S3 of SI). We note an increase in SASA values in Delta, suggesting a greater exposure of the 630-loop.

We propose that the structural rigidity of Omicron comes from higher inter- and intra-domain interactions of the S protein. Figures 4A and B represent the distribution of inter-protomer heavy atoms contacts (Figure 4A) and H-bonds (Figure 4B) between protomer-A and its neighboring protomer-B in the S1 domain. We observe a higher number of H-bonds and heavy-atom contacts in Omicron compared to Delta and D614G. We also calculated intra-protomer heavy-atom contacts between the S1 and S2 domains of protomer-A (Figure 4C). We noticed the same trend that Omicron forms more S1/S2 contacts than Delta and D614G. We also performed the same calculations for RBD-down protomers (Figure S4 of SI) and observed that Omicron forms more inter/intra domain contacts and H-bonds.

**Figure 4.**
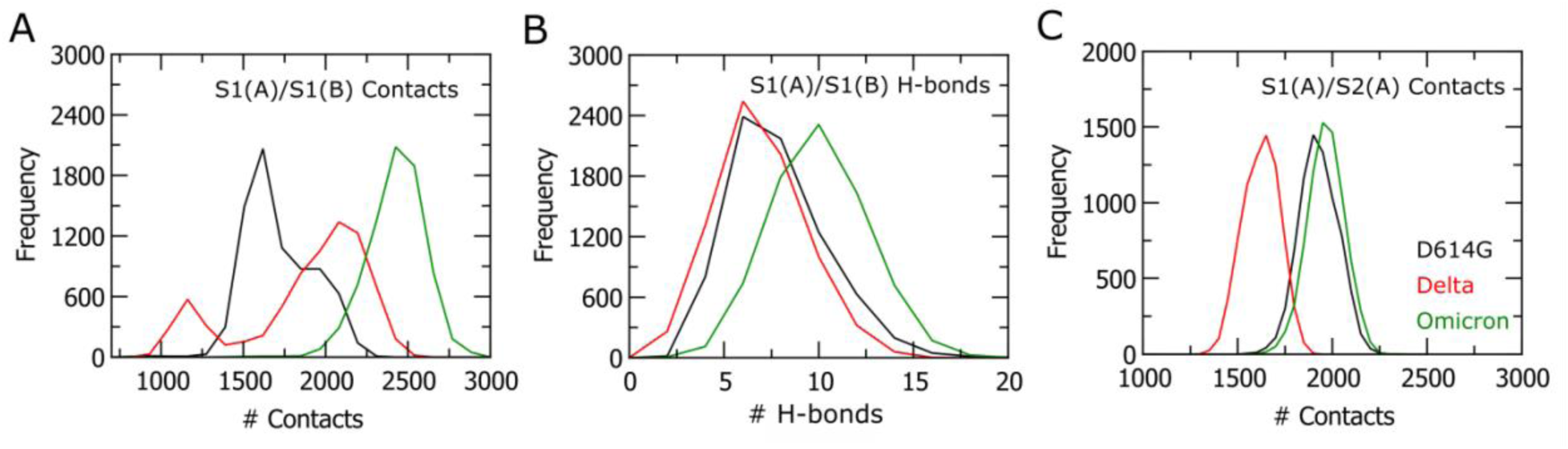
Inter/intra-domain contacts and H-bonds between protomer-A/protomer-B and protomer-A/protomer-A of the S1 domain. Panels (A) and (B) show the number of heavy-atom contacts and H-bonds distributions between protomer-A and protomer-B of S1, respectively. Panel (C) shows the number of heavy-atom contacts between S1 and S2 of protomer-A. Black, red, and green colors represent D614G, Delta, and Omicron.

To understand the contribution of each mutation to the structural rigidity/flexibility of S when the RBD is open, we calculated the number of heavy atom contacts of each residue of concern with the rest of the S protein (Figure 5). Here we considered mutated residues of Omicron (T547K, D614G, H655Y, N679K, P681H, N764K, D796Y, N856K, Q954H, N969K, L981F) and Delta (D614G, P681R, D950N) except the residues in the NTD and RBD domains and the calculations were performed for all the three variants. Overall, we observed that mutated residues in Omicron form a higher number of intra- and inter-protomer contacts (Figure 5A). We further focused on some of those residues to understand their local interactions. We saw that 614G residues in all three variants interact with 648G residues in the CTD2 domain of the same RBD-up protomer (Figure 5B). These interactions appear to inhibit premature shedding of S1, leading to the amplification of functional spikes. Several experimental studies have reported that D614G mutation enhances spike fitness and increases viral replication and infectivity.(*36, 40, 41*) A close-up look at the H655Y mutation in Omicron exhibits a *π* − *π* stacking interaction with the aromatic ring of the F643 residue in the CTD2 domain of the same protomer (Figure 5B). A cryo-EM study by Yurkovetskiy *et al.* reported that either D614G or H655Y mutation moved RBD towards an open conformation essential for ACE2 binding, thus enhancing the viral fitness(*42*). We found a cation-*π* inter-protomer interaction between N856K and D568 (Figure 5C). However, we did not observe any significant charged or polar interaction of L981F; rather, this residue was buried into the cluster of non-polar residues coming from HR1 and CH domains that could make impactful Van der Waals interactions (Figure 5D). We calculated relevant distances between those interacting residues as well, and the distributions demonstrate that the interactions are stable during the MD simulations (Figure S5 of SI).

**Figure 5.**
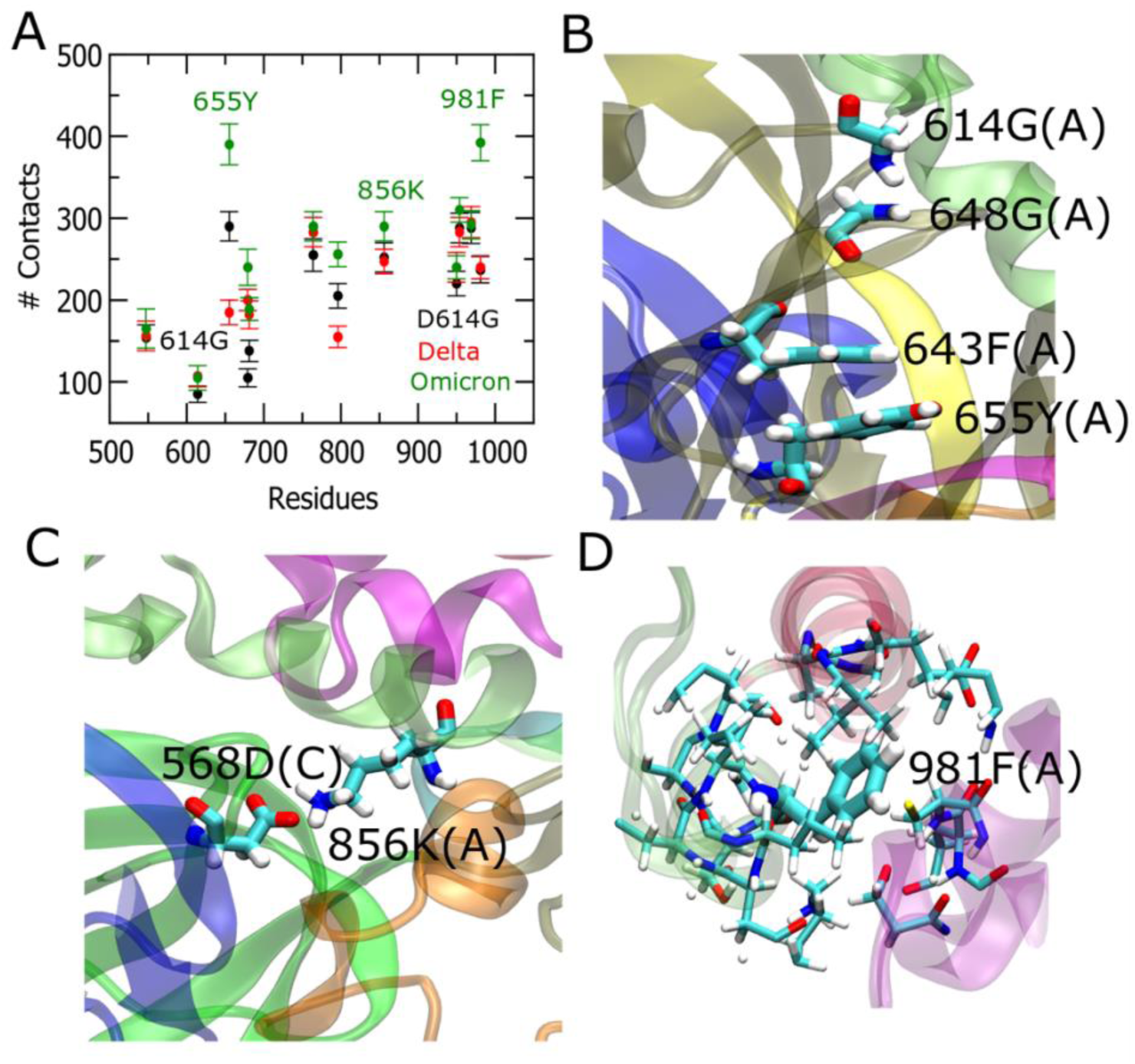
Contacts of each mutated residue with the remainder of the protein. Panel (A) shows heavy-atom contacts between mutated residues of protomer-A and the rest of the protein residues. It includes both inter and intra-domain contacts. Panels (B), (C) and (D) represent a close-up view of the region near the mutated residues of Omicron. D614G is forming an H-bond interaction with 648G of CTD2. A *π* − *π* stacking interaction was observed between 655Y and 643F (CTD2) of protomer-A. 856K (protomer-A) is forming a salt-bridge interaction with 568D of protomer-C. 981F is buried into a cluster of non-polar residues of HR1 and CH domains.

### P681R mutation enhances the fusogenicity in Delta compared to the P681H mutation in Omicron

The thermal fluctuations in the S1 domain are initiated by ACE2 binding and propagate to the S2 domain, especially into the fusion core, to instigate the fusion mechanisms. The junction between the S1 and S2 domains consists of a loop region (residues from 680 to 689) (Black circle in Figure 1B). We note mutations of P681 residues to ARG and PHE for Delta and Omicron in this loop region. Both ARG and PHE are large residues compared to PRO; therefore, these mutations introduce a steric hindrance in the loop. Figure 6A illustrates the distribution of the radius of gyration (R_g_) of the S1/S2 loop, which determines the unfolding of the loop. In this study, we observed that Delta exhibits a larger peak at higher R_g_ values (0.67 nm) compared to Omicron (0.55 nm). Additionally, a secondary peak was observed at 0.55 nm and 0.61 nm for delta and omicron variants, respectively. However, D614G shows a narrow distribution with a peak at 0.50 nm. These results indicate the highest amount of unfolding of the loop occurs for Delta, followed by Omicron and D614G. The presence of two consecutive bulky and positively changed residues (R681 and R682) in Delta creates a steric hindrance in the loop, resulting in the unfolding of the loop. Figure 6B depicts the loop structure of Delta and the locations of two ARG residues (R681 and R682). A close-up inspection of the superimposed structures of Omicron and Delta shows Delta has a more disordered loop where two ARG groups are facing opposite directions. This unfolding of the loop can initiate S1 shedding and facilitate the activation of S2 core, which is essential for viral membrane fusion(*43, 44*). A recent study by Alona *et al.*(*45*) revealed from the live virus plaque formation assays that the P681H mutation in Omicron modestly promotes cell fusion and syncytia formation, whereas the P681R mutation in Delta enhances fusogenicity and syncytia formation to a greater extent. When they introduced single point mutations of P681R in D614G and Omicron spike, they observed the significant restoration of fusogenicity. In contrast, when they mutated R681 to PRO in Delta, it notably reduced its fusion activity. Our results help to explain the critical mechanisms of P681R mutation in Delta for the enhancement of membrane fusion potential.

**Figure 6.**
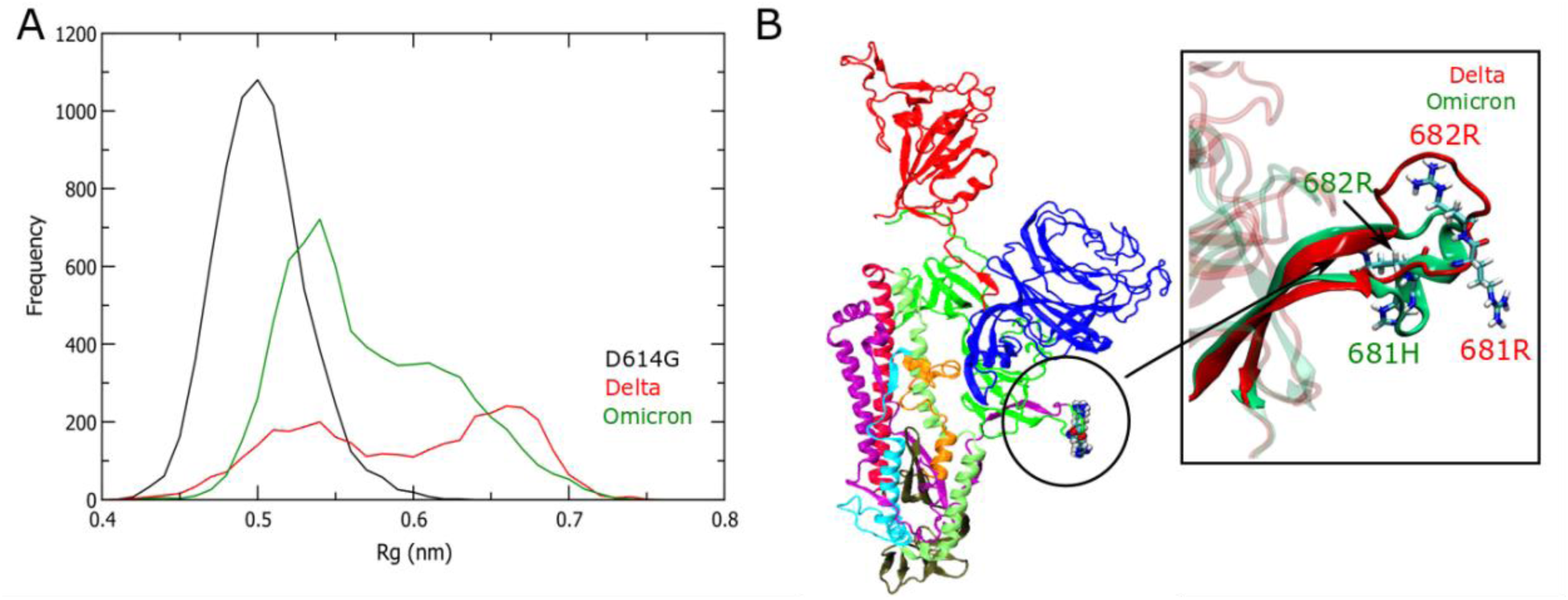
The radius of gyration plot and the loop at the S1/S2 junction of spike protomer-A. (A) The distribution of radius of gyration (Rg) values at the loop of S1/S2 junction (residue 680-689) of S protein. Black, red, and green colors represent D614G, Delta, and Omicron, respectively. Panel (B) shows the loop in the S1/S2 junction (black circle) in the Delta. Two ARG residues in the loop regions are highlighted to show the steric clash and unfolding of the loop. The zoomed picture shows the superimposed structure of the loop of Delta (red) and Omicron (green).

### Increased thermostability in Omicron inhibits pre- to post-fusion transition compared to Delta

During pre- to post-fusion transition, the S2 core undergoes complex conformational changes where a ’U-shape’ structure between HR1 and CH transforms into a single continuous helix(*16, 43*). The structures are shown in Figures 7A, B, and C. Here, we considered the ’U-shape’ structure as the bent conformation. From our RMSF calculations (Figure 3), we observed an increased fluctuation at the junction of HR1 and CH helices (HR1/CH hinge, residues 982-989) in the RBD-up protomer of Delta. To investigate the impact of this enhanced fluctuation in the conformational changes at the HR1/CH hinge, we calculated free energy (potential of mean force) based on backbone torsion tICA, taking eight residues (residues 982-989) at the HR1/CH hinge (Figure 7E, F, and G). We found minima for the bent conformations in all three variants (Figure 7E, F, and G). Only Delta shows a second minimum at higher tIC1 values, which implies a slightly stretched hinge structure (Figure 7F). The superimposition of the bent and stretched structures is shown in a box in Figure 7F. We took two residues (residues 983 and 987) in the hinge region and calculated C*α* distances between them for all three variants. Figure 7D shows a single peak close to 0.64 nm for D614G and 0.65 nm for Omicron, while Delta exhibits two peaks at 0.65 nm and 0.74 nm. These results again validate the presence of a stretched HR1/CH hinge conformation in Delta. The distribution of the dihedral angle in the HR1/CH hinge region (Figure S6A in SI) shows two distinct distributions in Delta in protomer A. For the RBD-down protomer, we observed only one distribution of dihedral angle at the hinge region (Figure S6B in SI) for all three variants.

**Figure 7.**
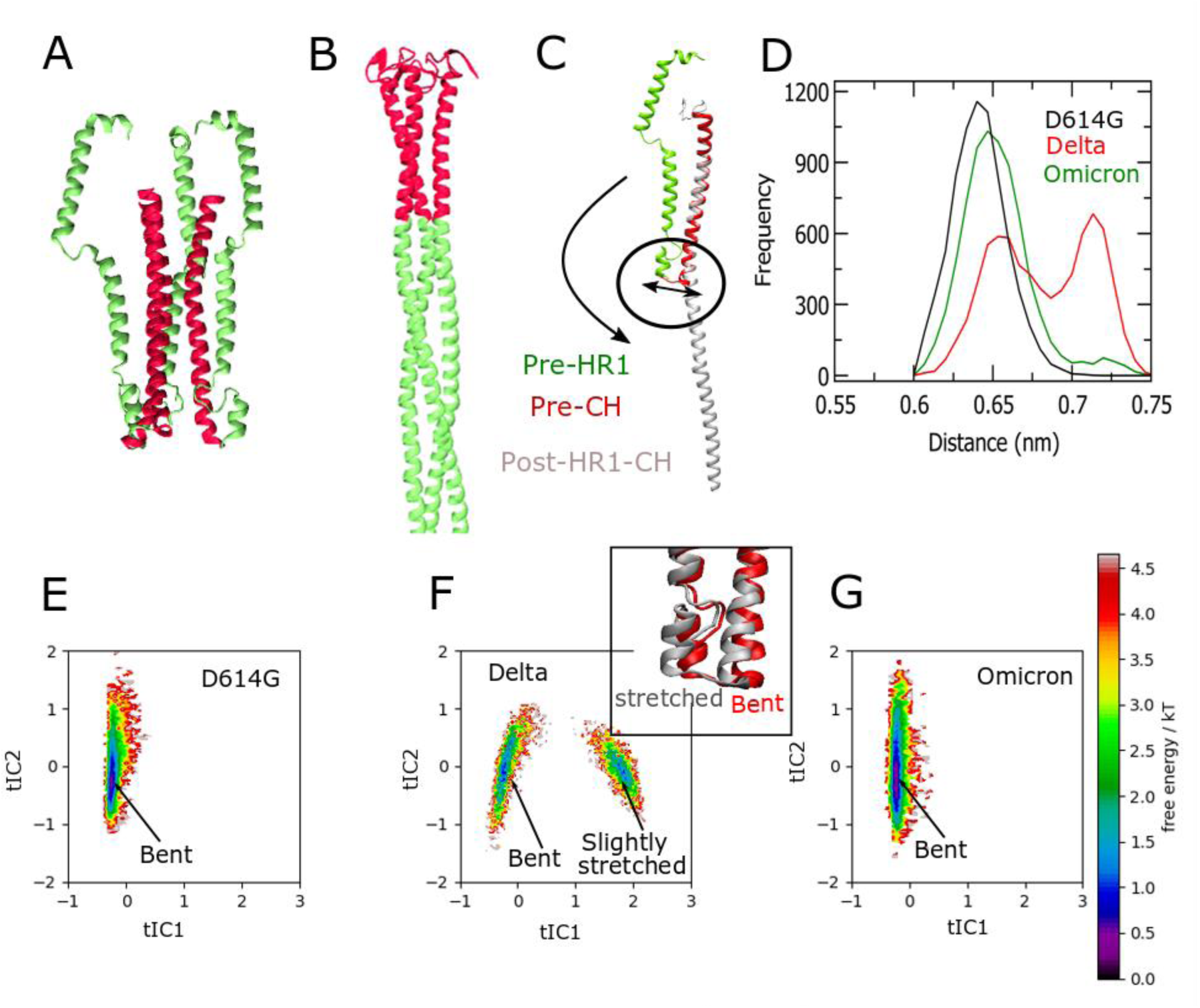
Pre- to post-fusion conformational changes of HR1 and CH domains and corresponding conformational sampling in tICA free energy plots. Panels (A) and (B) show pre- and post-fusion states of HR1 (green) and CH (red) domains of the spike protein. (C) Superimposition of pre- and post-fusion states of HR1 and CH domains. (D) End-to-end distance between two C*α* atoms (983 and 987) at the HR1/CH hinge. Panels (E), (F) and (G) show free energy plots of backbone torsional tICA of the HR1 and CH linker (the black circle in (C)). Delta has two minima at the bent and slightly stretched conformations. A close-up view is shown in the top right of Panel (F). D614G and Omicron exhibit a single minimum with bent conformation.

For further analysis, we considered two channels consisting of three HR1 helices (green helices in Figure 8E) and three CH helices (red helices in Figure 8E). During the pre- to post-fusion transition, the CH core remains intact(*43*), whereas the HR1 core alters its conformation from a bend to a straight structure (Figure 7C). The possible stepwise changes could be: (i) breaking the contacts between HR1 and CH, then (ii) swelling of the HR1 core, followed by (iii) straitening of the HR1/CH hinge. We calculated the minimum pore radius vs. the number of time frames and mean pore radius vs. distance along the central axis (z-axis) for three variants to explore the swelling of the HR1 channels. We observed the lowest values for minimum pore radius and mean pore radius in Omicron, while Delta shows the highest values for both (Figure 8A and B). The D614G variant exhibits values in between Delta and Omicron. We did not observe any significant changes in CH channels in the three variants (Figure 8C and D). The results indicate a slight swelling of the HR1 channel in Delta, which possibly enhances the transition in the HR1/CH hinge during pre- to post-fusion changes. We also noted a slight reduction in the number of contacts between HR1 and CH for Delta (Figure 8F). Recent experiments captured the intermediates when S proteins interact with ACE2(*46*). Braet *et al.* performed Amide hydrogen-deuterium exchange mass spectrometry on different SARS-CoV-2 variants and observed that Omicron shows a decreased exchange at the trimeric stalk, suggesting an increased stability of the stalk region(*47*). A recent study by Song *et al.* captured S-mediated virus-virus membrane fusion(*48*). They proposed a six-step membrane fusion model where dimpling is a state when the S2 core swells after S1 shedding. The observed swelling of the S2 core and stretching of the HR1/CH hinge, as observed in our simulations, point to an amplification of the cell-cell fusion process in Delta.

**Figure 8.**
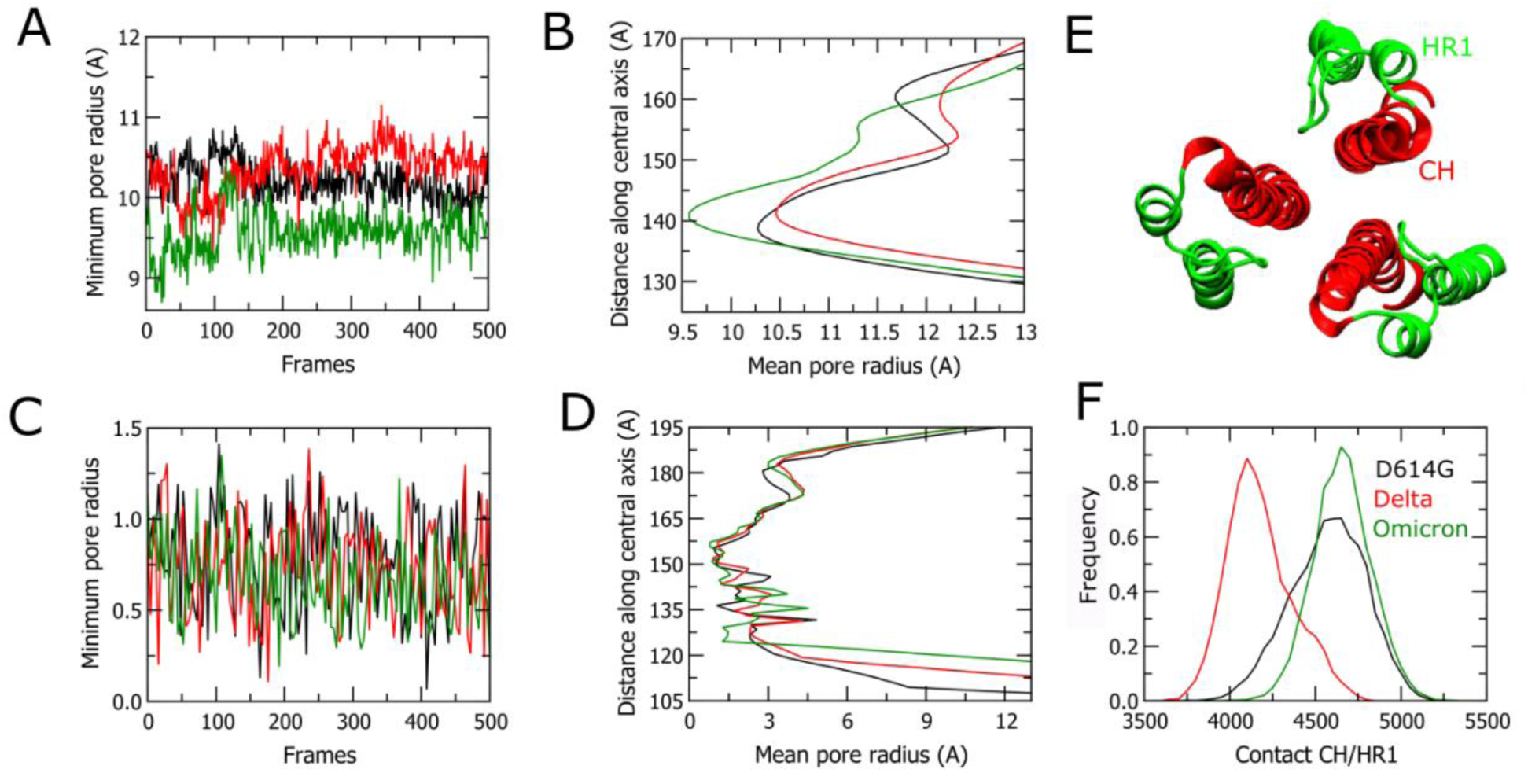
Calculation of minimum and mean pore radius of HR1 and CH helices. Panels (A) and (C) show the minimum pore radius vs. time frames plot for HR1 and CH helix channels, respectively. Panels (B) and (D) show mean pore radius vs. distance along the central axis for HR1 and CH helix channels, respectively. Black, red, and green colors indicate D614G, Delta, and Omicron, respectively. Panel (E) shows the HR1 (green) and CH (red) helix channels in a pre-fusion state. Panel (F) is a distribution plot of the number of all-atom contacts between HR1 and CH of protomer-A.

Our results further suggest that increased thermal fluctuation and reduced thermal stability in Delta lower the barrier for the activation of the S2 fusion core. This higher thermal fluctuation is initiated by ACE2 binding at RBD. The mutation P681R promotes S1/S2 cleavage and aids the activation of the fusion core. Omicron has a higher binding affinity towards ACE2 because of the charged mutations at the RBD; however, the mutations in the other subdomains of S1 and S2 generate a rigid and less flexible structure. Although the P681H mutation unfolds the loop at the S1/S2 junction to some extent, it still cannot compensate for the rigidity of the structure, which leads to a higher energy barrier for the pre- to post-fusion transition. D614G lacks a certain degree of affinity for ACE2. Furthermore, P681 reduces S1/S2 cleavage. Hence, D614G loses the competition of host cell membrane fusion even if RBD binding enhances the swing motion and thermal fluctuation in S1.

## Conclusions

In this study we have performed all-atom MD simulations of SARS-CoV-2 trimeric S protein in a complex with the ACE2 receptor for three SARS-CoV-2 variants, D614G, Delta, and Omicron. We provided a detailed analysis of the conformational dynamics and the role of the mutations in the propagation of the ACE2 binding effect from the RBD to the S2 fusion core. A large number of charged mutations reduces the structural flexibility of the Omicron S protein. A number of studies have reported that the Omicron RBD has a stronger binding affinity towards the receptor ACE2, which is a potential reason for its very high infectivity(*17, 20, 49*). It can easily spread from one person to another, which defines Omicron as a concerning variant of SARS-CoV-2. However, other experimental studies have suggested that the Delta variant shows higher viral loads in lung epithelial cells than Omicron(*50, 51*). Omicron appears to stay more in the nasal pathway and can only less readily infect lung cells. These results indicate an increased infectivity and a reduced severity for Omicron compared to Delta. Our results provide possible mechanisms for the contrasting roles of Delta and Omicron in infectivity and severity. Here, we also considered the original D614G variant to compare with the other two variants and to provide a more systematic study.

We observed that ACE2 binding at the RBD enhances the thermal fluctuations and structural flexibility of the S protein. The spike mutations play separate roles for Delta and Omicron to control the effect of the thermal fluctuations. Delta has three critical mutations, L452R and T478K at RBD and P681R at S1/S2 loop along with D614G(*52*). Besides RBD/ACE2 binding, P618R plays a key role in enhancing its fitness for membrane fusion(*53, 54*). Our results also show an unfolding at the S1/S2 loop due to the steric hindrance of P681R mutations in Delta. This unfolding enhances S1/S2 shedding and eases the activation of fusion core, ultimately promoting the complex conformational changes in the HR1 and CH domains during the pre- to post-fusion transition. D614G does not show any significant unfolding at the S1/S2 loop compared to Delta and Omicron.

Omicron S has 30 mutations, including D614G, and 15 of them are located in the RBD domains(*16*). Many of the mutations in Omicron appear to not be critical for viral fitness. However, the higher accumulation of positive charge at the surface of the S protein enhances the inter and intra-domain interactions, producing a relatively rigid structural conformation and reducing the thermal fluctuations. Omicron shows a moderate unfolding of the S1/S2 loop due to the P681H mutation.

We observed a small swelling of the HR1 channel at the fusion core for Delta. tICA free energy calculations based on backbone torsion of the HR1/CH hinge provide two minima for bent and slightly stretched conformations in Delta. Omicron and D614G show a single minimum with a bent conformation. We propose that increased thermal fluctuations reduce the transition barrier in the HR1/CH hinge of the Delta variant, thus expediting the pre- to post-fusion transition during viral membrane fusion. Therefore, Delta should need a smaller number of ACE2 to activate its fusion core than Omicron and D614G. It has been reported that lung cells have less ACE2 than the nasal path(*33*). Our study thus provides a possible explanation for why Delta shows higher viral infection in lung cells than Omicron.

In summary, this comprehensive all-atom MD study has yielded valuable information for understanding the molecular mechanisms by which the effect of ACE2 binding allosterically activates the fusion core in the three main SARS-CoV-2 variants so far.

## Methods

### System preparation and simulation details

The glycosylated structure of one RBD-up for D614G, Delta, and Omicron was taken from PDB IDs: 7KRQ(*36*), 7SBL(*52*), and 7TO4(*16*), respectively. Table S1 of SI shows the mutations in different locations of the S protein. To model the S trimer with the ACE2 complex, we used the RBD-ACE2 crystal structure (PDB 6M0J)(*55*) as an initial model for ACE2. We aligned the RBD-up domain of each spike protein with the RBD domain of the RBD-ACE2 complex structure. The initial glycosylated systems of S/ACE2 were prepared using CHARMM-GUI(*56–58*). We did not use any lipid membrane explicitly for our simulations. However, to consider the effect of the membrane, we restrained the position of the backbone atoms at the end segment of TM domains (residues residue 1150-1162). The simulations were performed with the GROMACS MD simulation package(*59*) using the CHARMM36m force field(*60*) and TIP3P water model(*61*). An initial energy minimization of the system was carried out following six-step protocols provided on CHARMM-GUI(*57*). We used an integration time step of 2 fs with periodic boundary conditions for the simulations. The simulation temperature was maintained at 310.15 K with a Nose-Hoover thermostat(*62, 63*) and a coupling time constant of 1.0 ps in GROMACS. The pressure was set at 1 bar with a Berendsen barostat(*64*) during initial relaxation. For the production runs, the Parrinello-Rahman barostat(*65*) was used semi-isotropically with a compressibility of 4.5×10^−5^ and a coupling time constant of 5.0 ps. For the nonbonded interactions, a switching function between 1.0 and 1.2 nm was used. The long-range electrostatics were computed using Particle Mesh Ewald(*66*). We used the LINCS algorithm to constrain hydrogen bonds(*67*). For each system, the simulations were performed for 3 μs with two replicas on the Frontera and Anton2 supercomputers (see Acknowledgements) and the Midway2 cluster (Research Computing Center at the University of Chicago).

### Analysis

For analysis of the MD trajectory results, we used the GROMACS tools(*59*), VMD scripts(*68*), and Python scripts, and MSMbuilder, MDAnalysis, and PyEMMA libraries(*69, 70*). The RMSF per residue, heavy-atom contacts, H-bonds, and R_g_ were calculated using GROMACS tools. All the heavy-atom and all-atom contacts were calculated by taking a 0.6 nm cut-off distance. The PCA was calculated using Python MDAnalysis libraries. Minimum pore radius vs. time frames and mean pore radius vs. distance along the central axis were calculated using Python MDAnalysis libraries. For these calculations, we stored data every 6 ns to reduce the computational cost. Protein structures were prepared using VMD and PyMOL(*68*). The tICA free energies were calculated using Python PyEMMA libraries. Our unbiased trajectories of the HR1/CH hinge region were used for the tICA calculation. The backbone dihedral angle of the hinge region was used as a feature space for the tICA analysis.

## Acknowledgments

This work was supported by the National Institutes of Health (NIH) grant R01AI169896. Computational resources were provided by the Frontera Supercomputer at the Texas Advanced Computer Center (TACC) funded by the NSF (OAC-1818253) and the Anton 2 machine funded by NIH (R01GM116961) at the Pittsburgh Super Computing Center (PSC). Some initial equilibrations of the systems were performed on Midway2 at the Research Computing Center (RCC) at the University of Chicago.

## Author Contributions

M.D and G.A.V designed, performed, and analyzed molecular dynamics simulations. M.D. and G.A.V wrote the paper. G.A.V. secured funding and supervised research.

## Notes

### Competing Interest Statement

The authors have declared no competing interest.

