## Supplemental Information for "Divergent spike mutations impact the activation of the fusion core in Delta and Omicron variants of SARS-CoV-2"

**This PDF file includes:**

Table S1

Figures S1-S6

Movies S1-S3

**Table S1. Mutations in the spike protein of D614G, delta, and omicron variants**

| D614G | Delta | Omicron |
|---|---|---|
| D614G | T19R, R158G, L452R, T478K, P681R, D950N | A67V, T95I, G142D, L212I, G339D, S371L, S373P, S375F, K417N, N440K, G446S, A477N, T478K, E484A, Q493K, G496S, Q498R, N501Y, Y505H, T547K, D614G, H655Y, N679K, P681H, N764K, D796Y, N856K, Q954H, N969K, L981F |

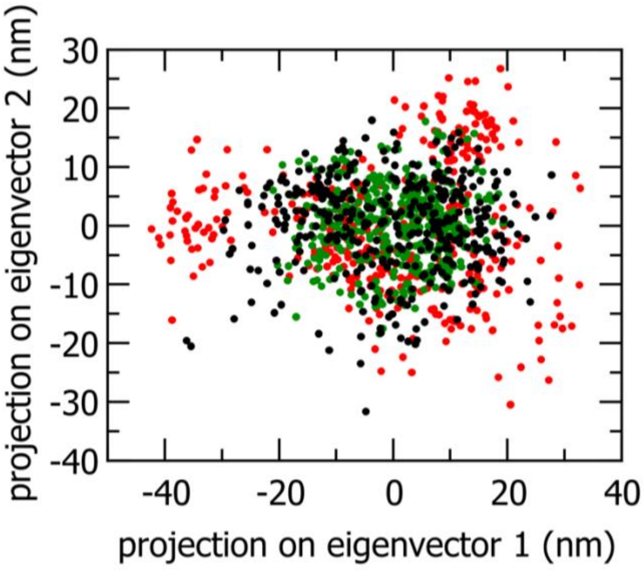


**Figure S1.** Projections of eigenvector-1 and eigenvector-2 calculated over C$\alpha$ coordinates of ACE2 bound spike. The black, red, and green colors represent D614G, delta, and omicron variants, respectively.

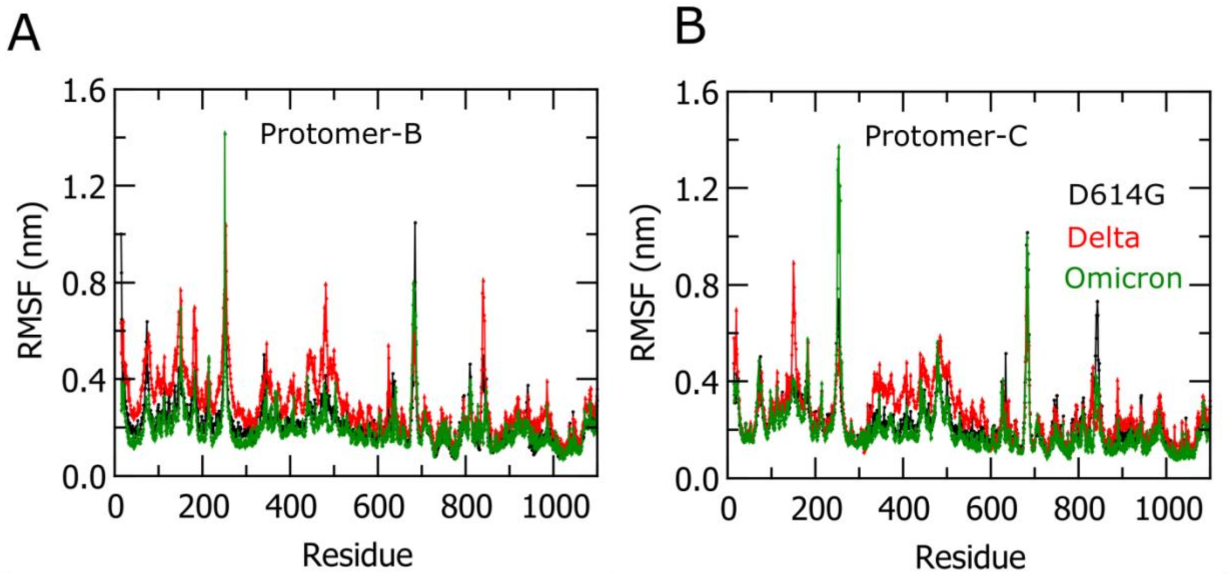


**Figure S2.** RMSF plots for RBD-down protomers. Panels (A) and (B) show protomers B and C respectively. D614G, Delta, and Omicron are represented in black, red and green colors, respectively.

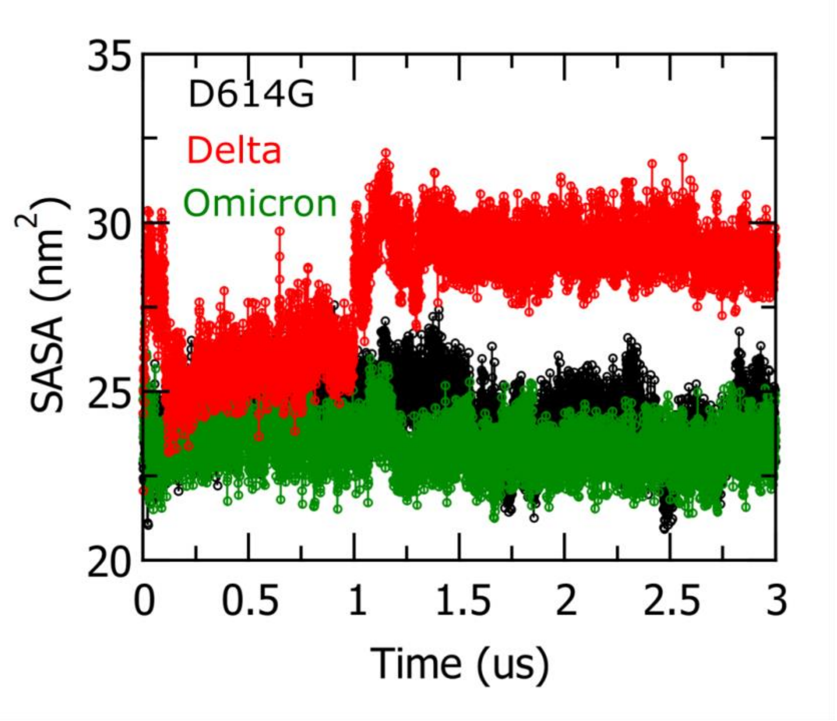


**Figure S3.** SASA vs. simulation time plot of 630-loop in D614G (black), delta (red), and omicron (green).

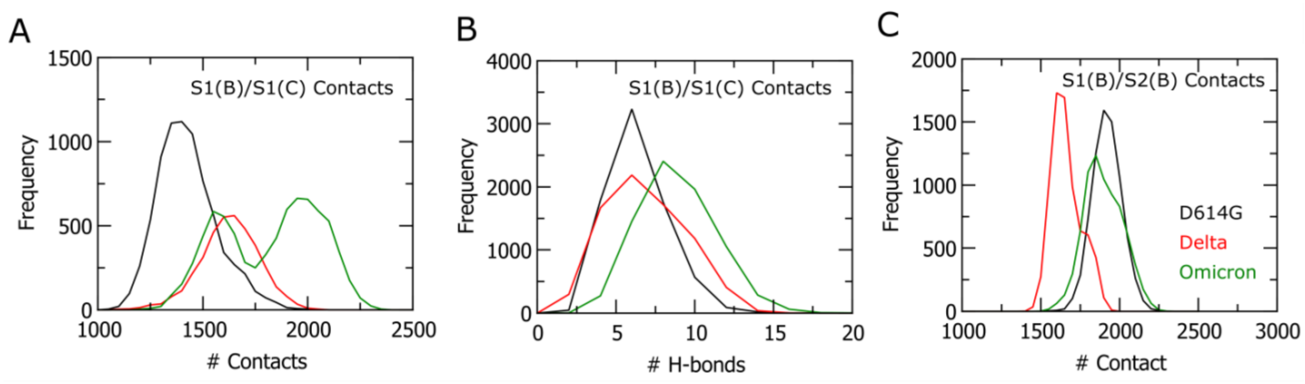


**Figure S4**. Inter/intra-domain contacts and H-bonds between protomer-B/protomer-C and protomer-B/protomer-B of S1 domain. Panels (A) and (B) show the number of heavy-atom contacts and H-bonds distributions between protomer-B and protomer-C of S1 respectively. Panel (C) shows the number of heavy-atom contacts between S1 and S2 of protomer-A. The black, red, and green colors represent D614G, delta and omicron.

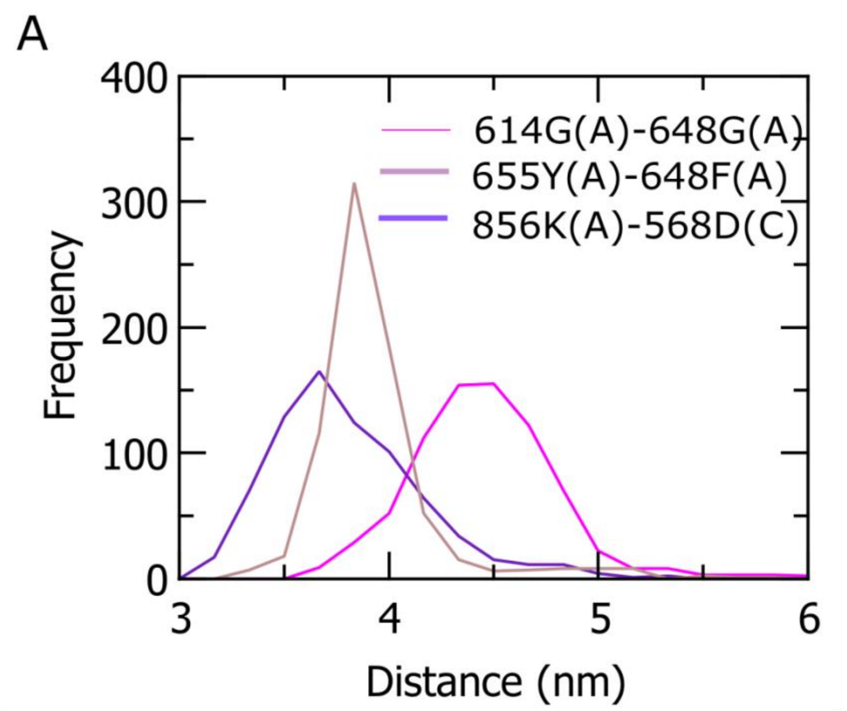


**Figure S5.** Distribution plot of distances of specific contacts between 614G(A) and 648((A) (magenta), 655Y(A) and 648F(A) (brown), 856(A) and 568(B) (violet).

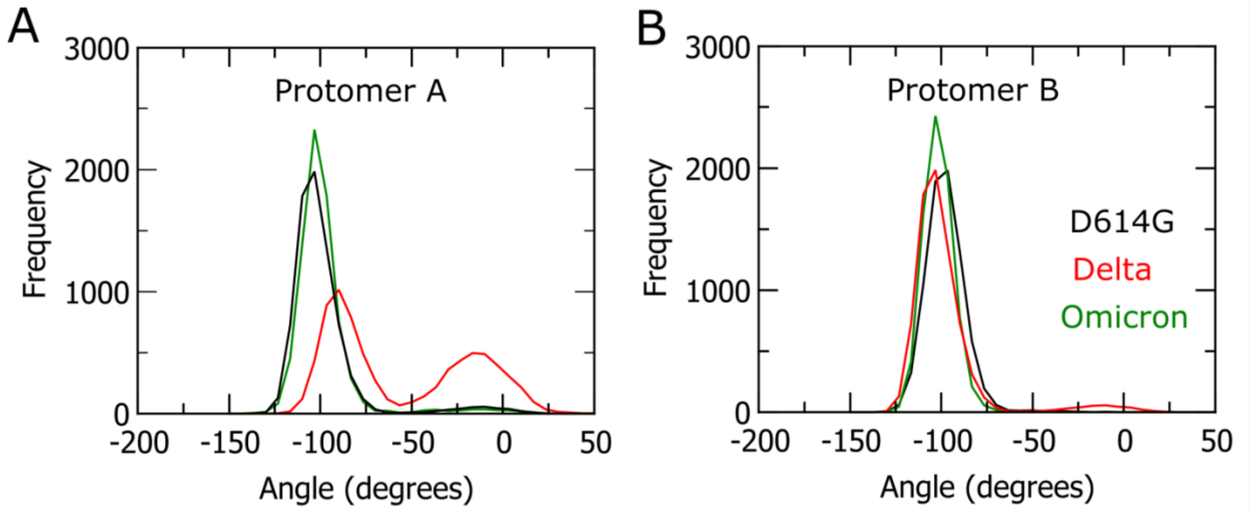


**Figure S6.** Distribution of dihedral angle at the HR1/CH hinge in protomer-A and B for all three variants.

**Movie S1-S3.** The first three principal motions of the RBD-ACE2 structure. Movie S1 corresponds to the swing motion of ACE2 bound RBD approaching and leaving its neighbor down RBD. Movie S2 describes a swing motion of ACE2-RBD towards its down and up states. Movie S3 depicts a motion toward the neighboring NTD domains.
